# Personalized anti-cancer drug combination prediction by an Integrated Multi-level Network

**DOI:** 10.1101/2020.05.12.092239

**Authors:** Fangyoumin Feng, Zhengtao Zhang, Guohui Ding, Lijian Hui, Yixue Li, Hong Li

## Abstract

Anti-cancer drug combination is an effective solution to improve treatment efficacy and overcome resistance. Here we propose a network-based method (DComboNet) to prioritize the candidate drug combinations. The level one model is to predict generalized anti-cancer drug combination effectiveness and level two model is to predict personalized drug combinations. By integrating drugs, genes, pathways and their associations, DComboNet achieves better performance than previous methods, with high AUC value of around 0.8. The level two model performs better than level one model by introducing cancer sample specific transcriptome data into network construction. DComboNet is further applied on finding combinable drugs for sorafenib in hepatocellular cancer, and the results are verified with literatures and cell line experiments. More importantly, three potential mechanism modes of combinations were inferred based on network analysis. In summary, DComboNet is valuable for prioritizing drug combination and the network model may facilitate the understanding of the combination mechanisms.

## Background

Cancer is the leading life-threatening disease across the world with more than eighteen million new diagnosed cancer cases and 9.6 million death in 2018 [1]. Due to the genetic and phenotypic heterogeneity of cancer, conventional anti-cancer monotherapies could not reach the expectation of clinical outcome. Unavoidable resistance and side effects induced by some monotherapies require effort on exploring more effective treatment strategies. Therefore, combining anti-cancer medicines has become a feasible alternative because of their advantages on sensitizing cancer response, modulating multiple biological progresses or pathways and reducing side effects[2]. Till 2015, only 49 anti-cancer combinatorial chemotherapies have been approved by FDA [3]. To discover more drug combinations, several high-throughput drug combination screening on cancer cell lines have been established which allow hundreds of drug pairs being tested in short time [4]. However, experimental screening all anti-cancer drug pairs exhaustively is impractical. Thus, in-silico discovery of potential drug combinations is considered as a reasonable way.

Two major strategies are considered for constructing more precise prediction models, that is to predict whether drugs can combine to achieve synergism and if the combinations can combine in a certain disease context. To address the former questions, methods like Zhao’s integrated drug-drug similarities including drug indication, drug ATC code, drug target proteins and drug side effects to predict effective drug combinations[5]. With the accumulation of cancer sample/cell lines transcriptome data and the understanding of molecular mechanism between drug and cancer, drug combination prediction in the context of cancer sample has gradually become the main direction. Databases like CCLE and LINCS released drug treated cancer cell line transcriptome data offer a solid base to support the construction of a cancer-specific dynamic network which reflect the real drug function [6-8]. Dialogue for Reverse Engineering Assessments and Methods consortium (DREAM) launched a worldwide open challenge in 2014 for drug synergy/combination prediction aimed at developing prediction models based on the integration of multilevel data [9]. Among the 31 submitted models, DIGRE, the top one algorithm, predict drug synergy based on modelling drug combination induced transcriptome changes from monotherapy perturbations[10]. SynGen predict combinable drug pairs which work complementary towards master regulators who induce cell death or inhibiting cell status activation [9]. Following the challenge, Cao et al, proposed a well performed model compared with other methods based on semi-supervised learning called RACS[11]. It integrated seven features from drug targeting networks and two filtering parameters from transcriptomic profiles and predicted potential drug combination based on the similarities to positive dataset. Though the features in RACS and other models were more focusing on local similarity between drug targets or the function of targets, the integration of multi-level drug related information and the combination of targeting network and drug treatment transcriptome profile provide a new notion of building a dynamic disease network interpreted by drug treatment. With the accumulation of high-throughput drug screening data, several supervised models have been built based on large high-throughput drug synergy screen dataset, like 39 drugs for 38 cancer cell lines provided by O’Neil [12] and 710 drug combinations across 85 cancer cell lines in drug synergy prediction DREAM challenge [13]. Algorithms like DMIS, NAD and Y Guan, performed in top three position in DREAM challenge, predict drug synergism on cell lines via multi-dimensional feature extraction and machine learning methods[13]. These algorithms showed good performance on the data set provided by DREAM[13]. Later on, deep learning model Deepsynergy used similar strategy achieve good performance[14]. However, these supervised learning algorithms tend to have high dependence on a large number of cell line experimental data to achieve good predictions on the corresponding cell lines. Furthermore, the algorithms are usually difficult to apply on other cell lines than the modelling set. When tested on the O’Neil data set, the performance of Y Guan, DMIS, NAD are all dropped[13]. Although performed well on cross-validation on modelling dataset, Deepsynergy did not verify in external dataset[14].

The complexity of drug mechanism on real cancer context is still a main obstacle in combination prediction. Constructing a heterogeneous network is an applicable solution for integrating multilevel information and modeling different biological systems[15, 16]. Han’s method mapped drug on gene expression profile weighted PPI network via drug-target associations to predict drug-drug interactions [17]. WNS method evaluated drug synergy in pathway-pathway interaction network [18]. DrugComboRanker discovered potential drug combination by identifying drug-drug associations in target networks[19]. Barabasi’s synergy prediction is based on measuring the distance between drug modules and disease modules discovered from the gene network between drug targets and disease related genes[20]. In addition to use in predicting synergistic drug combinations, network-based approaches may help infer potential mechanism between combinable drug pairs via the construction of biological network[15]. However, these network-based methods based mainly on drug and their known direct target genes, even with the integration of pre-treatment transcriptome data, they did not show enough power to predict sample-specific drug combinations.

Assuming pharmacologically similar and functionally related drugs tend to combine together, we proposed a computational method called DComboNet to predict the anti-cancer drug combination. The DComboNet level one model constructs a generalized heterogeneous network integrating drug-drug, drug-gene, drug-pathway, gene-gene and pathway-pathway associations. Drugs that can be combined with the drug seed are predicted according to their global similarity in the network. The level two model constructs a cancer sample specific network to predict personalized drug combination. DComboNet was evaluated using cross validation, independent test and experimental validation. DComboNet outperformed the previous methods. Additionally, DComboNet provides clues for the potential mechanisms of drug combinations.

## Results

### Characteristics of know anti-cancer drug combinations

We collected 218 known anti-cancer drug combinations that involved in 157 drugs. There were three types of drugs: 67 standard chemotherapy (C), 57 targeted cancer therapy (T) and 33 other kind of drugs (O) (**Fig. 1a)**. The effects and anti-tumor mechanisms of these three types drugs are distinctive. Standard chemotherapy acts on both normal and cancerous cells via their cytotoxic function; targeted cancer therapies are deliberately chosen or designed to interact with their specific target or targets with a cytostatic mechanism; other drugs may help control cancer related complications or alleviate on side effects caused by anti-cancer medicine. The combinations within and between three types can all be seen in known anti-cancer drug combination (**Fig. 1b**).

**Figure 1.**
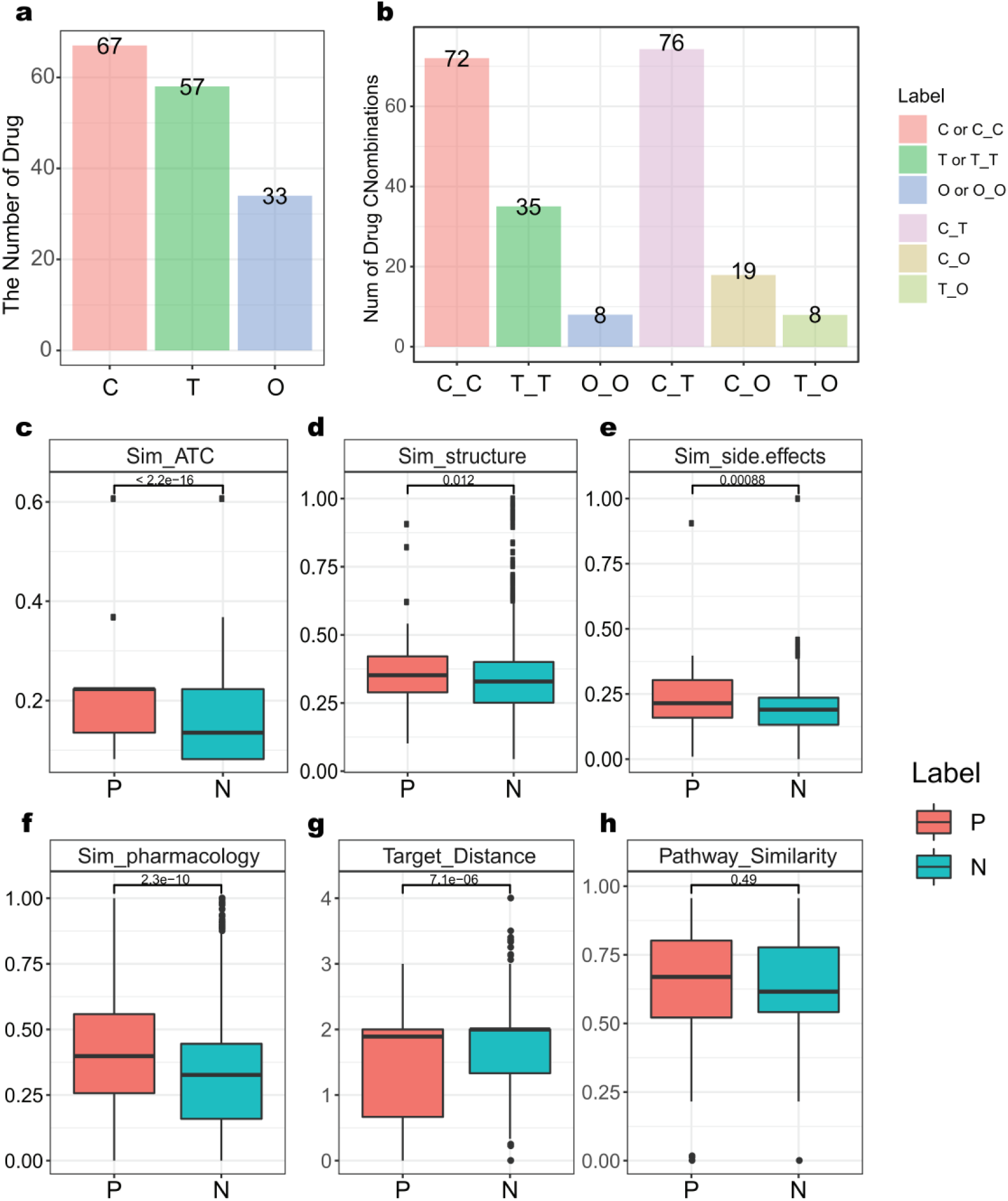
Features of known anti-cancer drug combinations. **a** The distribution of drug types. C, T and O are standard chemotherapy, targeted cancer therapy and other cancer-related drugs, respectively. **b** The distribution of drug combinations by drug types. **c-e** Comparison of ATC code similarity, chemical structural similarity and side-effect similarity between known drug combinations (P) and unlabeled drug pairs (N); **f-h** Comparison of the integrated pharmacological similarity, target distance and pathway similarity between known anti-cancer drug combinations (P) and unlabeled drug pairs (N).

The mechanisms of drug combinations can be partially explained by pharmacological similarity, topological associations of drug targeted genes and functional pathways[5, 11, 18]. To understand the contribution of these features to anti-cancer drug combinations, we generated 9017 unlabeled drug pairs by randomly pairing the drugs in the positive dataset and then compared unlabeled pairs with known combinations. The ATC code, chemical structure fingerprints and side effects were used to calculated the similarity between drug pairs respectively (**Supplementary methods**), and then combined into an integrated pharmacological similarity. All the single and integrated pharmacological similarity were higher in known combinations than in unlabeled pairs (**Fig.1c-f**). Target distance was the average distance between two target gene sets on the background gene-gene interaction network. Drugs in known combinations had shorter target distance than in unlabeled pairs (**Fig. 1g**). Pathway similarity between drugs was implemented via computing the average shortest distance between pathways, and if drugs co-regulate same pathway, using the shortest distance between their targeted genes to represent their pathway similarity. (**Supplementary methods**). The pathway similarity of drugs in known combinations is also higher than randomized pairs, though not significant (**Fig1h**). Therefore, we thought integrating both drug pharmacological and functional associations may help predict combinable drug pairs and reveal the potential mechanisms.

### Workflow of cancer drug combination network (DComboNet)

The concept of DComboNet is to abstract pharmacological and functional relationships between drugs into a heterogeneous cancer drug combination network **(Fig 2a**). DComboNet consists of five subnetworks: drug-drug association network (*N*_*DD*_), drug-gene association network (*N*_*DG*_), gene-gene association network (*N*_*GG*_), drug-pathway association network (*N*_*DP*_) and pathway-pathway association network (*N*_*PP*_).

**Figure 2.**
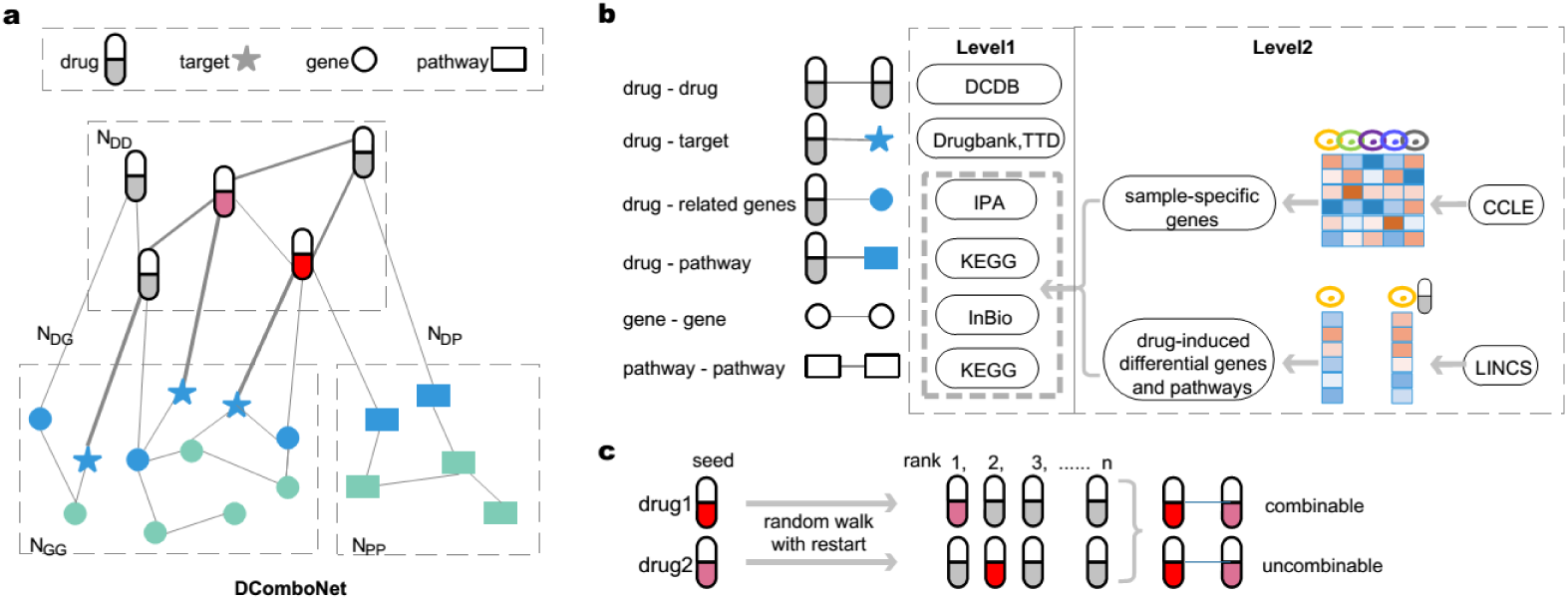
Workflow of drug combination prediction model (DComboNet). **a** Construction of heterogeneous cancer drug combination network (DComboNet). The network contains five sub-networks, N_DD_ indicates drug-drug association network, N_DG_ indicates drug-gene association network, N_GG_ indicates gene-gene association network, N_DP_ indicates drug-pathway association network and N_PP_ indicates pathway-pathway association network. **b** Source and methods for generating network edges. Level one indicates the generalized model and level two indicates cancer sample specific model. **c** Method of ranking drug pairs.

DComboNet contains two levels of models (**Fig. 2b**). Level one is a general model that predict the potential drug combinations. It was established without considering individual heterogeneity. Edges were generated from multiple databases and the weights of edges were assigned based on the edge types (seeing **Methods**). Level one model may be not precise enough for specific cancer type or individual sample. Introducing transcriptome data into drug combination prediction can enhance the precision of the prediction for certain cancer type [10]. Therefore, level two model utilized transcriptome data to reconstruct networks and predict drug combinations for a specific cancer sample. Sample specific expressed genes were obtained by comparing the expression profile of this sample with other cancer samples. Drug induced differentially expressed genes and pathways were obtained by comparing the expression profiles before and after drug treatment.

After network construction, Random Walk with Restart method was applied to capture the global proximity between the given drug seed and candidate drugs in the network. For a drug pair, drug1 and drug2, take each of them as seed to calculate the global proximity between drug1 and drug2, respectively. Then a two-threshold strategy was used to integrate two ranks and classify the drug pair into combinable, uncombinable and intermediate (**Supplementary methods**).

### Performance of level one model

Leave-one-combination-out cross validation (LOCOCV) was used to evaluate the performance of level one model. Firstly, we compared the prediction performance of using drug-drug network alone and using the integration of multiple different networks. DComboNet integrated five subnetworks and obtained the best performance (**Fig.s1c**). The AUC of DComboNet is 0.797 and the true positive rate (TPR) is 63.24%. Secondly, we compared the prediction results of different drug types (**Fig. 3 a-b**). Standard Chemotherapy combinations (C-C) performed well with AUC equals to 0.816. All 68 real drug combinations within this category were ranked in top 50%, of which 51 known combinations were predicted to as combinable. Targeted therapy drug combinations (T-T) also achieved high accuracy with 17 out of 23 known combinations were predicted correctly (TPR = 76%). Due to the lack of pharmacological and functional associations between standard chemotherapy and targeted therapy, the accuracy of C-T combinations is slightly less powerful (TPR = 53.52%). Lastly, we compared DComboNet with a previous published algorithm, RACS preliminary model, which also predict without using transcriptome data [11]. DComboNet outperformance RACS which has AUC equal of 0.548 and true positive rate 46.0% (**Fig. 3c**).

**Figure 3.**
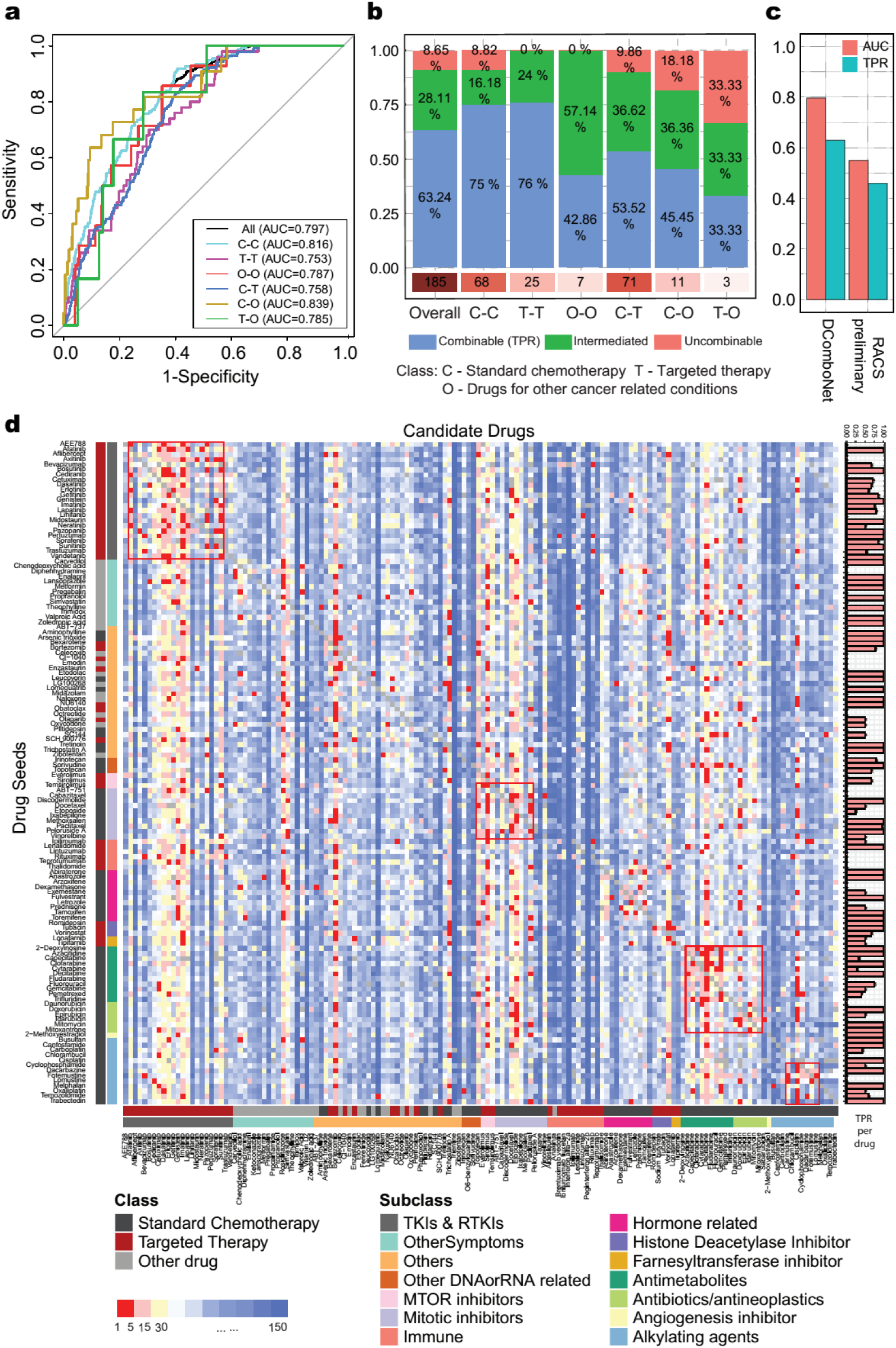
Performance of the level one model. **a** ROC plots and AUC values based on different types of drug combinations. **b** The percentage of predicted as combinable, uncombinable and intermediate in known drug combinations with respect to different drug combination types. **c** Comparison of performance between DComboNet and RACS preliminary. **d** The result of Level one model DComboNet. Heatmap was sorted in two directions according to drug subclasses (**Table s1**). Each row represents a drug seed. Color in the plot shows the rank of similarity between drug seed and other drugs. The bar plot on the right shows the successfully predicted drug pairs in known combinations (TPR for each drug seed).

Furthermore, we used DComboNet to predict the combination potentially of all drug pairs. (**Fig. 3d**). In order to better analyze the prediction result, we further categorize drugs into 12 subtypes based on their mechanism of action. We found drugs within the same subtype are more likely to be recommended as combinable drugs because of their relevant functions, such as inhibitors of tyrosine kinases and their receptors, drugs that interfere mitotic or target on microtubule (red box in **Fig. 3d)**.

### Performance of level two model

Performance of the DComboNet level two model was first evaluated using hepatocellular carcinoma cell line HepG2 and breast adenocarcinoma cell line MCF7. Take HepG2 as an example, gene expression profiles between HEPG2 and other cancer cell lines were compared to obtain specifically expressed genes in HepG2; gene expression profiles of HepG2 before and after monotherapy were compared to generate the differentially expressed genes (DEGs). HepG2 specific network was constructed by added 632 HepG2 specifically expressed genes, 78913 drug-DEG, 1652 drug-DEpathway associations and the corresponding gene-gene, gene-drug and gene-pathway associations. Potential drug combinations were predicted by level two model based on HepG2 specific network. The AUC of DComboNet level two model was 0.844 for HepG2, rising significantly compared with AUC of level one model (AUC = 0.787, **Fig. 4a**). The prediction on breast cancer cell lines MCF7 was also improved with the AUC rose from 0.764 to 0.795 (**Fig. 4b**).

**Figure 4.**
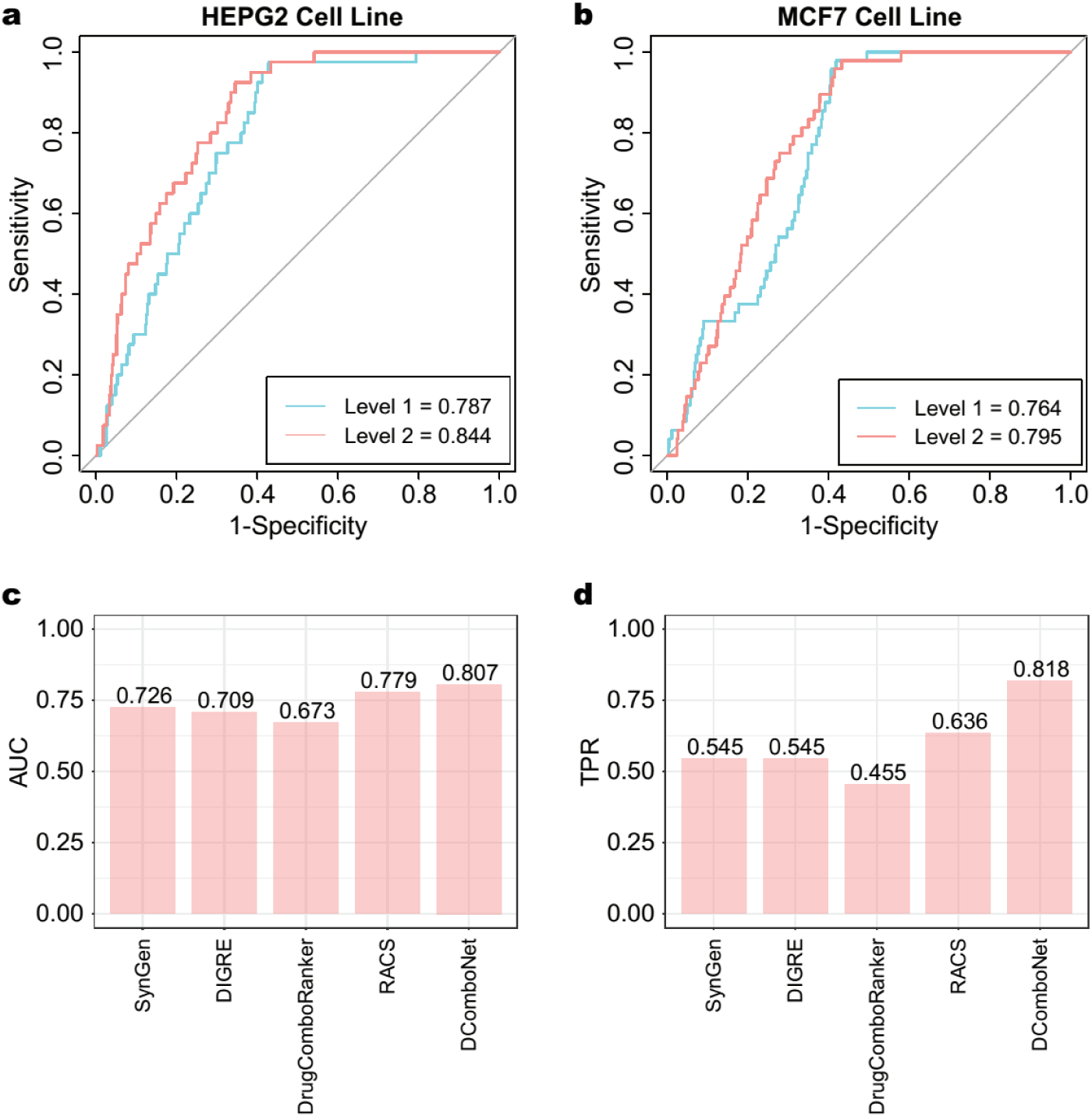
Performance of cancer sample specific drug combination prediction model. **a)** and **b)** ROC curves of model on hepatocellular carcinoma cell line, HEPG2, and breast cancer cell line, MCF7. In each plot, red and blue curves indicate the ROC curves of level two model and level one model, and the number in legend indicate the AUC values. **c-d)** Method comparison between DComboNet level two model and other prediction algorithms (SynGen, DIGRE, DrugComboRanker, RACS) using the OCI-LY3 dataset.

Some drug combinations, especially standard chemotherapy that directly act on DNA/RNA, cannot be correctly predicted in the level one model. By adding drug perturbated transcriptome change, the effects of these drugs on cancer cells can be reflected through changes in a series of genes or pathways instead of only their target genes, therefore correct prediction may be obtained. For example, the combination of capecitabine and docetaxel are both standard chemotherapies with a broad anti-cancer effect[21]. Although they show relatively high pharmacological drug similarity (*sim*_*DD*_ (capecitabine, docetaxel) = 0.695), the target genes and the biological functions between capecitabine and docetaxel are distinctive (target distance is 2, pathway similarity is 0.45) [22, 23]. This combination failed at predicting as combinable pair in level one model, but was successfully predicted in level two model after reconstructing cancer sample specific network.

We further compared DComboNet level two model and other four drug combination prediction algorithms (SynGen[9], DIGRE[10], DrugComboRanker [19], and RACS[11]) which also used the change of transcriptome profiles before and after monotherapy treatments. All of these algorithms were evaluated using the drug synergy screening dataset (OCI-LY3). The overall performance of DComboNet outperformed other algorithms, especially more powerful when predicting the combinable pairs (**Fig 4c-d**). DComboNet achieves 0.807 AUC and 81.8% true positive rate. Among 11 real synergy pairs, 9 pairs were successfully predicted by DComboNet while 7, 6, 6 and 5 pairs were successfully predicted by RACS, SynGen, DIGRE and DrugComboRanker, respectively (**Fig 4d**).

## Case study: HepG2 - sorafenib

Hepatocellular carcinoma (HCC) is the fourth leading cause of cancer death with only few approved agents as first line treatment, such as sorafenib[24-26]. However, most patients will develop sorafenib resistance eventually which include multiple biological pathways. Therefore, it is critical to find potential drug combination to improve the efficacy of single sorafenib treatment. We predicted combinable drugs for sorafenib treatment through DComboNet level two model and validated the predictions through literature investigation and in-vitro experiments. Using ‘Sorafenib’ as the drug seed, 5 out of 6 known combinations were predicted correctly. In the rest 26 newly predicted combinable drug pairs, 16 of them have been reported to be synergistic either in HCC models (8 pairs) or in other cancers (8 pairs) in previous literatures (**Fig.5a**).

In addition to predicting the propensity of drug combinations, DComboNet can also rank genes and pathways in the network according to the proximity relative to drug seed. Thus, analyzing the overall results may be helpful in inferring the possible mechanism of drug combinations. Among the drugs predicted as combinable candidates for sorafenib, we found three potential mechanism modes for effective combination (**Fig 5b-d**).

**Figure 5.**
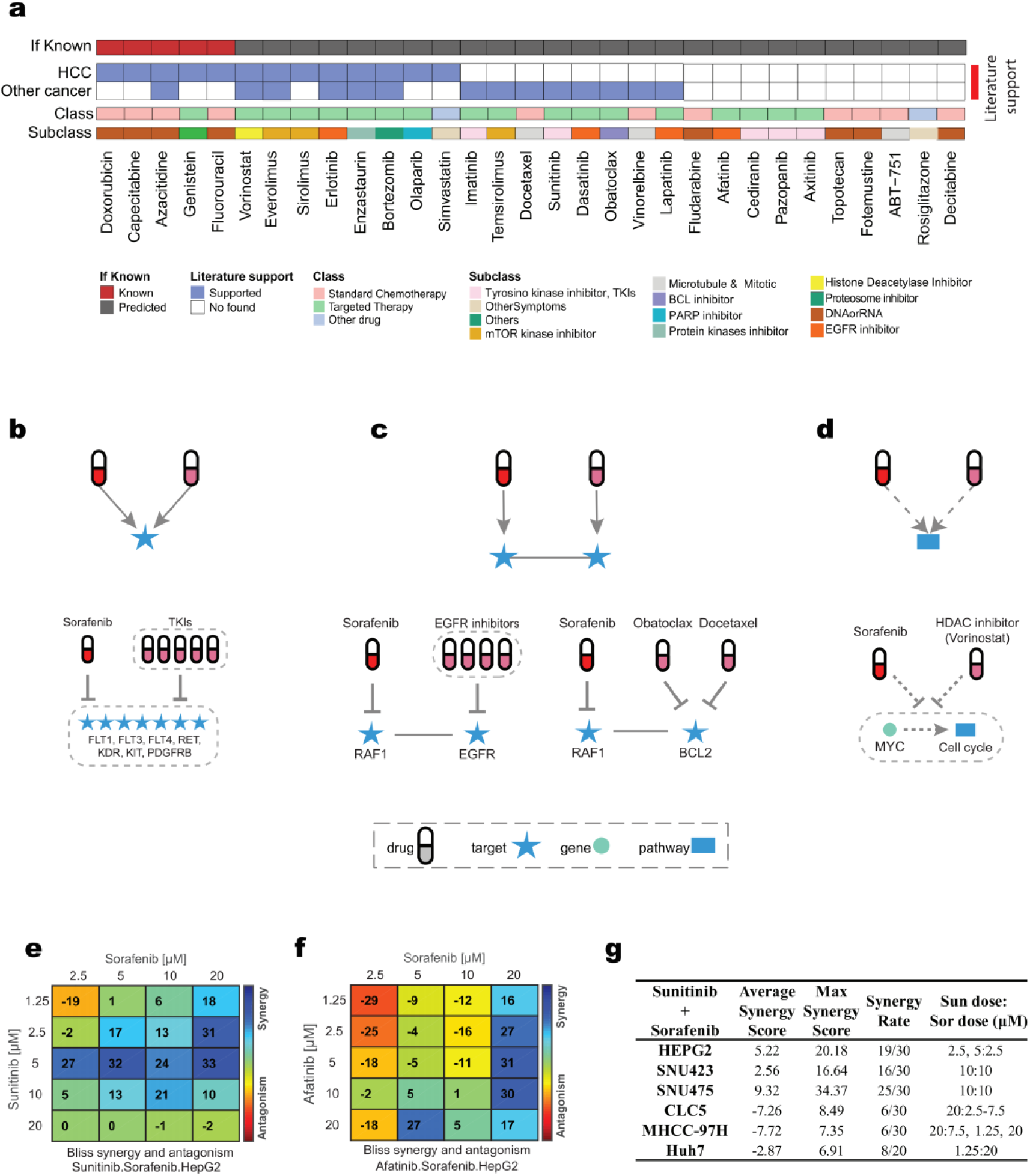
Predictions and validation for Sorafenib on Hepatocellular carcinoma cell line, HEPG2 and the hypothetical mechanisms inferred. **a)** Overall prediction results for Sorafenib on HEPG2 cell line. The first line denotes if the predicted combinable drugs belong to known drug combination for Hepatocellular carcinoma (HCC); the second and third lines denote if predictions have literature supports for HCC or other cancer types; the fourth and fifth lines show the classification information of predicted drugs. **b-d)** indicate schematic diagrams of three potential combination mechanism modes for sorafenib case study. The upper part of **b)** shows the schematic diagram for drugs targeting on the same genes to achieve synergy, and the bottom shows the example “sorafenib-other tyrosine kinase inhibitors (TKIs)” matching this mode. The upper part of **c)** shows the mode that the interaction between drugs’ target genes lead to synergy, and the bottom part shows two examples (sorafenib and EGFR inhibitor and sorafenib and BCL2 inhibitor). The upper part of **d** shows the synergy may through the regulation of cancer-related genes other than target genes, and the matching examples (Sorafenib and HDAC inhibitors). In figure **b-d)**, the capsule shape nodes represent drugs, dark red corresponds to sorafenib, and light red corresponds to the predicted combinable drugs; blue stars represents target genes of drug; green round dots represents other genes in gene network; and blue rectangles represent pathways. **e-f)** Experimental validation results for sorafenib combined with sunitinib and afatinib, respectively. Each of the heatmaps shows the synergy score calculated by the Bliss method for each dose points. The color bar of heatmap shows the score range from synergy (blue) to antagonism (red). **g)** The summary table of experimental synergy screening for sorafenib and sunitinib in six hepatocellular carcinoma cell lines (HEPG2, SNU475, SNU472, Huh-7, CLC5 and MHCC-97H) with multiple dose combinations.

The first mechanism is that two drugs shared same target genes (**Fig.5b**). Among the prediction results, several multiple tyrosine kinase inhibitors (TKIs) show strong tendency to be combined with sorafenib. Imatinib, cediranib, dasatinib, sunitinib and pazopanib shared 6 targets (FLT1, FLT3, FLT4, KDR, KIT, PDGFRB) with sorafenib (Fig.5b). The combination between TKIs and sorafenib may work on cancer-related genes or functions through compensatory way to avoid single TKI resistance and further improve efficacy on cancer patients[27, 28]. For example, sorafenib can block the function of genes related to imatinib resistance in HCC treatment [29].

The second mechanism is that two drugs may achieve synergy through the regulatory relationships between their target genes (**Fig.5c**). Three epidermal growth factor receptor (EGFR) inhibitors, erlotinib, afatinib and lapatinib[22, 30] were predicted as candidates with combination potential with sorafenib. DComboNet showed that EGFR inhibitors connect to sorafenib through the ‘EGFR-RAF1’ link (**Fig.5c).** EGFR is an upstream signal receptor Ras pathway while RAF1 acts as a signal transduction mediate with RAS/RAF/MEK/ERK signaling pathway [31, 32]. Inhibiting EGFR can help sensitize the efficacy of RAF inhibitor (e.g. sorafenib) in hepatocellular cancer cell lines [33] and the synergism between RAF inhibitor sorafenib and EGFR inhibitors have also been observed in multiple cancer types [34-36]. Another example is the predicted combination of BCL2 inhibitor (docetaxel and obatoclax) and sorafenib. BCL2 inhibitor is connected with sorafenib via the association of their target genes “BCL2-RAF1” in the network **(Fig.5c)**. Over-expression of BCL2 and RAF1 may lead to sorafenib resistance, which can be altered by inhibiting BCL2 in HCC cell lines [37, 38]. This indicates that coadministration of BCL inhibitor and sorafenib may improve treatment efficacy.

The third mechanism of drugs combination is that they co-regulate cancer-related genes, even there are no direct target gene associations involved **(Fig.5d).** Take histone deacetylase (HDACs) inhibitors vorinostat as an example (**Fig.5d**). Vorinostat itself plays an important anti-cancer role which inhibit cancer cell growth via blocking cell cycle [39]. In the potential mechanism network, we found sorafenib and vorinostat are linked together via down-regulating MYC (**Fig.5d**). HDACi has been reported to help acetylate c-MYC and promote apoptosis in AML [40]. The sorafenib-vorinostat combination may coregulate multiple pathways related to cancer cell cycle and apoptosis to achieve synergism [41].

Based on these drug combination mechanisms, we selected two drugs (sunitinib and afatinib) to further verify the predicted combination with sorafenib in HCC. Sunitinib shared 7 target genes with sorafenib (**Fig.5b**), and their combination showed synergistic efficacy in renal cell carcinoma [42]. However, there are no similar studies in HCC. Therefore, we performed the combination experiments of sorafenib and sunitinib using HCC cell line HepG2. 14 of the 20 dose combinations showed synergy, and the most synergistic dose combination was when sunitinib was 5 μM and sorafenib was 20 μM (**Fig.5e)**. Furthermore, we verified a completely new prediction result, the combination of sorafenib and afatinib. Afatinib is an EGFR inhibitor, which may achieve synergy with sorafenib through the regulatory relationships between their target genes (**Fig.5c**). Experiments in HepG2 showed 9 synergistic points at different dose, indicating that sorafenib and afatinib is combinable (**Fig.5f**).

Additionally, the combination of sorafenib and sunitinib was further tested using another five HCC cell lines (**Fig.5g)**. SNU475 and SNU432 also showed synergy in experimental screening especially strong synergy at multiple doses, while synergistic effect on Huh7, CLC5 and MHCC-97H cell lines only occurred in few dose points. This reflects the heterogeneous response of cancer cell lines to the same treatment. It is necessary to make individualized drug combination prediction. If the expression profile of individual cancer sample is available, the DComboNet level two model could obtain personalized prediction results by utilizing sample specific network.

## Discussion

Discover efficient drug combination under the highly complex and heterogeneous cancer system is difficult and time-consuming on wet-lab synergistic drug screening whereas in-silico drug combination prediction has become a critical approach in preclinical research. Based on the comprehension of anti-cancer drug mechanism and the accumulation of cancer related data, we developed a two-level prediction model DComboNet. Level one model can be used to predict anti-cancer drug combination in a more general manner, whereas level two model is capable to achieve cancer sample specific drug combination prediction by integrating sample specific expressed genes, differentially expressed genes and biological pathways after drug treatment into the ‘drug-gene/pathway’ network.

DComboNet has several advantages: 1) DComboNet utilizes the complex multi-layer heterogeneous networks, which efficiently integrate multi-level data and provide more information to rank the combinable tendency from a holistic point of view. Therefore, DComboNet is possible to predict drug pairs that have a more intricate combination mechanisms other than the direct target gene association. 2) DComboNet contains two levels of models, which users can choose according to their aim and available data. Level two model has better prediction accuracy than level one, but requires the expression profiles of cancer sample before and after monotherapy treatment. 3) Compared to other algorithms using similar input data, DComboNet achieves higher true positive rate. 4) DComboNet provides drug-gene/pathway network between the predicted combinable drug pairs, which is helpful for understanding the potential mechanism of drug synergy.

We noted that certain drug combinations are usually poorly predicted, especially those with lower pharmacological similarities and less functionally relationship in gene or pathway network. Additionally, DComboNet ranks candidate drugs based on their global similarity with the seed drug, therefore it may have less power on predicting drug combinations with distinct mechanisms. With the accumulation of high-throughput drug screening data, we plan to combine DComboNet with supervised machine learning algorithms to improve the prediction performance. We also realized that the different response between cancer cell lines and patients under the same drug treatment still remains as a common obstacle for drug discovery transformation. Drug absorption, distribution, metabolism and excretion process cannot be well modelled under the context of cancer cell lines. Patient-derived mouse xenograft may serve as a better model than cell line for these, but drug screening on animal model like mouse need more effort and funding. Although there are many difficulties in the translation from basic research to application, computational prediction of drug combinations is fast and convenient as well as achieves much better accuracy than random. We anticipate that DComboNet could provide candidates for drug combination experiments and accelerate the discovery of new synergistic drugs.

## Methods

### Data collection

Known drug combinations were collected from DCDB 2.0 [43]. Only FDA approved drugs or drugs entering phase III or IV of clinical trial were kept in the subsequent analysis. Therapeutic information used to construct drug-drug association network included: drug ATC (Anatomical Therapeutic Chemical Classification System) codes[44] extracted from WHO Collaborating Centre for Drug Statistics Methodology (WHOCC) website, chemical structures downloaded from DrugBank[22, 45] and PubChem[46], drug side effect information collected from SIDER4[47]. Drug target proteins or genes were retrieved from Drugbank and Therapeutic Target Database (TTD)[48]; Drug related genes were obtained from IPA (Ingenuity® Pathway Analysis).

The human protein-protein interaction network were extracted from the scored InBio Map [49]. The interaction pairs with low score (score < 0.15) were removed. Cancer related genes were extracted from KEGG Cancer related pathways [50].

Gene expression profiles of drug perturbated cancer cell lines were downloaded from LINCS database[8] and DREAM challenge 2014[9]. LINCS database provided gene expression microarray data for 1127 cell lines treated by 41847 molecules. Drug treated hepatocellular carcinoma cell line HepG2 and breast cancer cell line MCF7 were extracted from LINCS. The pretreated gene expression data of HepG2 and MCF7 were downloaded from CCLE database[6].

### Level one model: Cancer Drug Combination Network (DComboNet)

Cancer Drug Combination Network (DComboNet) is based on a multi-level heterogenous network which contains five subnetworks, drug-drug association network (N_DD_), drug-gene association network (N_DG_) and gene-gene association network (N_GG_), drug-pathway association network (N_DP_) and pathway-pathway association network (N_PP_). The details of subnetwork construction are described in Supplement methods. Briefly speaking, N_DD_ obtained drugs from known drug combinations and their associations was weighted by pharmacological similarity integrated three kinds of drug similarity. For level one model, N_DG_ was constructed based on drug and target (D-T) associations, drug and drug-related gene (D-G) associations. N_GG_ integrated both cancers related genes extracted from KEGG cancer related pathway including ‘pathway in cancer’ and genes connected with drugs in N_DG_, and the associations between genes were extracted from inBio Map (V 2016_09_12) [49]. N_DP_ was constructed based on the association between drugs and their possible regulated pathways. N_PP_ was built on the hierarchy of KEGG provided in WNS method [18].

The network can be represented as an adjacency matrix 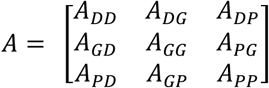, where *A*_*GD*_ and *A*_*PD*_ are transpose of *A*_*DG*_ and *A*_*DP*_. Given a certain drug, DComboNet recommends drugs with closest topological relationship as the combinable candidates. This global proximity between drugs can be captured via random walking with restart (RWR) algorithm. This algorithm originally designed to simulate a random walker walking on the network with certain initial probability corresponding network. For our task, we assigned the walk starts only from N_DD_. More specific, the random walker is assigned as a drug seed with an initial probability 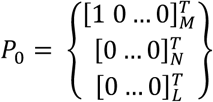, where M, N and L indicate the node number in N_DD_, N_GG_ and N_PP_, respectively. Walker will start from this drug seed node to traverse every node in network. At every step, the jumping happens from the current node to it direct neighbor(s) with a probability 1 − σ or returns to the drug node with a restart probability σ. The probability in t + 1 step can be represented as follow:

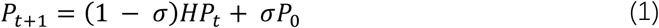

After several iterations, the probability will reach a steady state when the difference between P_t+1_ and P_t_ falls below 10^−10^. At this point, all nodes in the complex network have obtained global proximity relative to the drug seed which can be considered as the combination potential. After removing the known drug combinations, the rest candidate drugs can be ranked according to their potential.

In function (1), 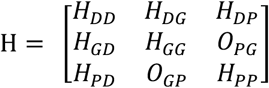 denotes the transition matrix which reflects different strategies for the walker to traverse the complex network.

The transition probability between drug 1 (*d*1) and drug 2 (*d*2) can then be described as:

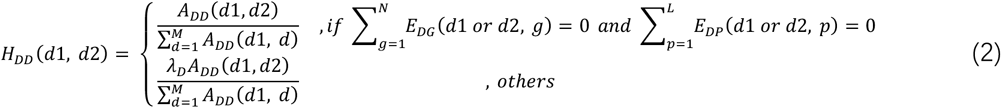

The jumping within drug network contains two different possibilities: if drug does not have any link with N_GG_ or N_PP_, the jump happens in N_DD_ with probability λ_D_ = 1; if *d*1 or *d*2 can be linked to any gene node (*g*) and/or pathway node (*p*), the potential jumping happens within N_DD_ with the probability λ _D_.

If drug can be linked to N_GG_, jumping from *d*1 to gene 1 (*g*1) may happen with the probability λ_DG_ and the transition probability can be described as:

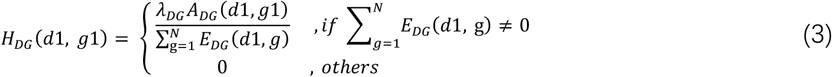

After the jumping fall in N_GG_, the transition probability from gene *g*1 to gene *g*2 in N_GG_ can be influenced by the existence of edges in N_GD_. Therefore, the transition probability within N_GG_ can be computed as:

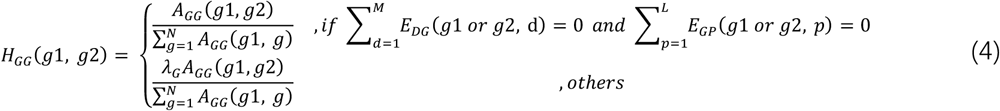

Similarly, if jump happens from N_GG_ back to N_DD_, transition probability between *g*2 and *d*2 can be described as:

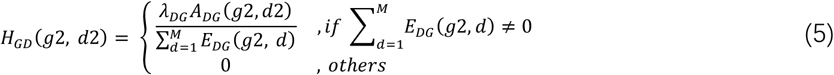

Similar to the jumping strategy through gene node, transition probability between *d*1 and pathway node *p*1 can be calculated as:

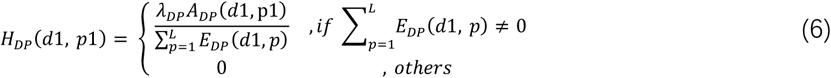

When the jump falls in N_PP_, we expected the next step can directly happen from N_PP_ back to N_DD_. More specific, if the edge(s) between drug and pathway exist, jump can only happen either within N_PP_ or between N_DP_. The calculation of transition probability within N_PP_ can be seen as follow:

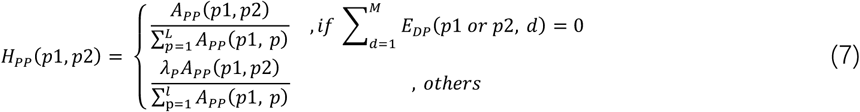

The jump from p2 back to d2 can then be described as:

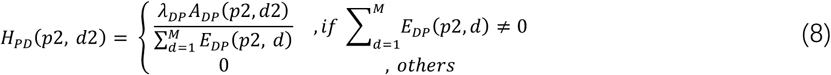

From the description of jumping strategy between d1 and d2, we can see that jumping probability λ are not independent. Within the homological subnetworks, λs are equal (*λ*_*D*_ = *λ*_*G*_ = *λ*_*P*_). The jumping probability in heterogenous subnetwork (such as N_DG_ and N_DP_) are influenced by those within homological subnetworks (*λ*_*DG*_ = 1 − *λ*_*D*_ and *λ*_*DP*_ = 1 − *λ*_*D*_).

To improve the accuracy of the model, global restart probability σ and jumping probabilities (λ_D_) were set to values from 0 to 1 and the optimal parameters were selected through cross-validation (supplement figure 1). Based on the tuning results, the default setting of λ_D_ is set as 0.5 to keep the balanced contribution of sub-networks, and σ is set as 0.7 which is also consistence with previous publications [51].

### Level two model: cancer-specific DComboNet

To predict sample specific drug combination, transcriptome data before and after drug perturbation were further integrated in the N_DG_ and N_GG_ as well as N_DP_ and N_PP_ to construct sample specific complex network.

The specifically expressed genes (*G*_*cancer*_) were selected with the criteria 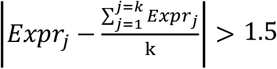 (that the fold change of gene expression between the specific cancer cell line and the average of expression value of all the other cell lines above 1.5). *G*_*cancer*_ were used to replace the nodes in *N*_*GG*_ for reconstruct cancer-specific gene-gene association network.

Cancer specific drug-gene and drug-pathway associations were added into the original drug-gene/pathway association network *N*_*DG*_ and *N*_*DP*_. These two associations were obtained by comparing drug treated gene expression data and DMSO. Differentially expressed genes were selected by functions lmFIt and eBayes in the Limma package (FDR < 0.1) [52]. Differential regulated pathways (DEpathway) were obtained by the GSVA algorithm (FDR <0.1) [53]. The edge weight of both drug-DEG were assigned as the fold changes of genes perturbated by drugs and drug-DEpathway were assigned as 1. Furthermore, these differentially expressed genes and their protein-protein interactions extracted from InBio Map were also added into *N*_*GG*_.

Similar to Level two model, a given drug was set to be the seed with initial probability *P*_0_ and iterated till the steady status of the probability of nodes. Then the candidate drugs will be ranked based on the final probability after removing the drug pairs in positive set.

### Cross validation and independent test

We conducted Leave-One-Drug-pair-Out Cross Validation (LODOCV) to assess the model performance. For each known drug combination, the edge weight was replaced by its integrated pharmacological similarity score and every drug in the combination will be used as drug seed to rank the rest of drugs in the network. Receiver operating characteristic (ROC) curves and the area under these curves (AUC) were also used to quantify the performance. To access the successfully predicted drug pairs and avoid the asymmetrical ranks, that is, the difference between the rank of B when A is used as drug seed and the rank of A when B is taken as seed, a two-threshold strategy was used to classify the drug pair into combinable, uncombinable and intermediate (Supplementary method).

To further verify the predictability and generalization of our level two model in independent dataset, we tested the model performance using the OCI-LY3 dataset [9]. Excess over Bliss (E.o.B.) and signal to noise ratio (s.n.r.) were calculated by the Bliss independent model [9]. Drug pairs were classified into combinable pairs (synergistic, E.o.B.>0 and s.n.r.>2) and uncombinable pairs (antagonistic and additive).

### Experimental validation

Potential combinations between sorafenib and sunitinib, sorafenib and afatinib were evaluated using liver cancer cell lines (Supplementary method). Cell viability matrices of each drug pair on the corresponding cell lines were used as the input data to calculate experimental synergy score with Bliss model provided by Combenefit software [54]. To reflect the degree of synergy, the maximum synergy score (Max Syn) and the synergy rate among all dose combinations was calculated.

### Code availability

DComboNet is implemented in R language and available at https://github.com/VeronicaFung/DComboNet.

## ACKNOWLEDGEMENTS

This work was supported by the National Natural Science Foundation of China 31771472), National Key Research and Development Project (2019YFC1315804), Chinese Academy of Sciences (ZDBS-SSW-DQC-02), SA-SIBS Scholarship Program, Shanghai Municipal Science and Technology Major Project (No.2018SHZDZX01), LCNBI and ZJLab.

## AUTHOR CONTRIBUTIONS

F.Y.M.F. and H.L. designed the study; F.Y.M.F. performed model construction and data analysis;

Z.T.Z. performed drug combination screening experiments; F.Y.M.F. wrote the manuscript; F.Y.M.F. developed the R package; Y.X.L. and H.L. supervised research and revised the manuscript.

