## Supplemental Method and Figures for "Personalized anti-cancer drug combination prediction by an Integrated Multi-level Network"

### Supplementary

#### Supplementary methods

##### 1. Three features used to calculate integrated pharmacological score.

**ATC code similarity**

The hierarchical structure of drug ATC (Anatomical Therapeutic Chemical Classification System) codes allows us to calculate drug indication-based similarity based on a probabilistic model [1]. Let $t_i$ $and$ $t_j$ represent the two ATC codes in the ATC hierarchical network, the similarity between two ATC codes ($t_i$ $and$ $t_j$) can then be calculated as:

$$sim_{ATC}(t_i, t_j) = w(t_i)w(t_j)exp\left(-\gamma d(t_i, t_j)\right) \quad (1)$$

where $d(t_i, t_j)$ denotes the shortest distance between $t_i$ $and$ $t_j$, $w(t_i)$ $and$ $w(t_j)$ represent the weights of the corresponding ATC codes, and are defined as the inverse of ATC-code frequencies, which means that more emphasis was put on specific codes rather than the general ones [2]. $\gamma$ is a pre-defined parameter (set to be 0.25 in this study).

According to WHOCC, ATC codes was assigned only to approved drugs and usually only one for each drug, even though drugs may have multiple indications. Therefore, ATC code similarity $sim_{ATC}(t_i, t_j)$ can be used as drug ATC code-based similarity $sim_{ATC}(A, B)$. However, some special cases like Prednisolone exist, which one drug may have more than one ATC codes. For this situation, the drug ATC similarity was considered as the average ATC code similarities between two drugs. For a pair has one or zero drug without ATC codes, their ATC code-based similarity score $sim_{ATC}(A, B) = 0$.

**Chemical fingerprint-based similarity**

The 2D chemical structures of drugs were downloaded from Pubchem. The binary vectors of chemical fingerprints were generated from chemical structure using software PaDEL[3]. Tanimoto index was used to calculate the similarity matrix[4]:

$$sim_{structure}(A, B) = \frac{\boldsymbol{v_A} \cdot \boldsymbol{v_B}}{\|\boldsymbol{v_A}\|^2 + \|\boldsymbol{v_B}\|^2 - \boldsymbol{v_A} \cdot v_B} \quad (2)$$

where $\boldsymbol{v_A}$ and $\boldsymbol{v_B}$ represent the chemical descriptor vectors of drug A and B. The similarities were calculated based on three kind of fingerprints included 1024 CDK fingerprints, 166 MACCS fingerprints and 881 PubChem fingerprints and averaged as the final chemical fingerprint-based similarity.

**Side effect-based similarity.**

Side effect information of drugs was integrated from SIDER4.0[5]. Then Jaccard coefficient was considered to compute the side effect-based similarity[6]:

$$sim_{sideeffect}(\mathrm{A}, \mathrm{B}) = \frac{\|S_A \cap S_B\|}{\|S_A \cup S_B\|} \quad (3)$$

where $S_A$ and $S_B$ represent the set of side effects of drug A and drug B respectively.

#### 2. Two topological associations of drug targeted genes and functional pathways

**Target distance**

Let M and N represent target gene sets belong to drug A and drug B separately, the similarity between drug a and b can then be defined as[7]:

$$\text{targetsim}(\text{a}, \text{b}) = \frac{\sum_{T_i \in M,\, T_j \in N} d(T_i, T_j)}{|M| \times |N|} \tag{4}$$

where $T_i$ and $T_j$ represents the targeted genes belong to gene set M and N respectively, $d(T_i, T_j)$ represents the shortest distance between two genes in the background gene-gene interaction network.

**Pathway similarity**

Let A and B represent two drugs in a combination, we first projected the drug to functional pathways through their targeted gene sets M and N to create pathway sets $P_A$ and $P_B$. If drug A and drug B correspond to different pathways, the pathway-based drug similarity was calculated as the average distance between pathways under the context of pathway-pathway interaction network derived from KEGG[8]:

$$\text{pathsim}(\text{A}, \text{B}) = e^{-\frac{\sum_{P_i \in P_A, P_j \in P_B} d(P_i, P_j)}{|P_A| \times |P_B|}} \tag{5}$$

where $d(P_i, P_j)$ represents the shortest distance between pathway i and j in the background pathway-pathway interaction network.

If drug A and drug B target the same pathway, i.e. $P_i = P_j$, the connection between A and B is considered as the average distance between targeted genes within the certain pathway:

$$\text{d}(P_i, P_j) = e^{-\frac{\sum_{T_i \in M, T_j \in N} d(T_i, T_j)}{|M| \times |N|}} \tag{6}$$

#### 3. Network construction in DComboNet

**Drug-drug association network**

The drug-drug association network ($\mathrm{N_{DD}}$) consisted of drugs obtained from known drug combinations and their relationships as edges. The weights of edges were defined by an integrated pharmacological similarity:

$$w_{DD}(\mathrm{a},\mathrm{b}) = \begin{cases} 1, & if\ E(a,b) \in M \\ 1 - \prod_{i=1}^{n}(1 - sim_i), & if E(a,b) \notin M \end{cases} \tag{7}$$

Where a and b represent two drug nodes in $\mathrm{N_{DD}}$; $\mathrm{E(a,b)}$ denotes the edge between a and b; M refers to the set of known drug combinations; $sim_i$ includes three kinds of drug similarity, drug indication-based similarity ($sim_{ATC}$), chemical structure fingerprint-based similarity ($sim_{structure}$) and drug side effect similarity ($sim_{sideeffect}$). The details of these drug similarities are further describing in **Supplement Methods**

**Drug-gene association network**

To better capture the functional interactions between drug pairs, drug-gene associations network ($\mathrm{N_{DG}}$) was constructed based on two kinds of associations, drug and target (D-T) associations, drug and drug-related gene (D-G) associations. The gene/protein names were remapped HGNC symbol[9] via biomaRt R package[10] and removed the duplicated association. Since target genes were considered as direct effects of drugs whereas drug related genes were more indirect effects, the weight ratios between D-T and D-G were set to 2:1.

**Gene-gene association network**

The gene-gene association network ($\mathrm{N_{GG}}$) integrated both cancers related genes extracted from KEGG cancer related pathway including 'pathway in cancer' and genes connected with drugs in $\mathrm{N_{DG}}$. Genes were mapped to official gene names with biomaRt and merged together as nodes of $\mathrm{N_{GG}}$. The associations between gene nodes were extracted from protein-protein interaction (PPI) network inBio Map (V 2016_09_12) [11] and the duplicated and loops of the network were removed. The edge weight in $\mathrm{N_{GG}}$ were set to be 1 if the association exists.

**Drug-pathway association network**

The drug-pathway association network ($\mathrm{N_{DP}}$) presents the relationship between drugs and their potential biological functions. The connection between drug and pathway is via a target-pathway mapping solution, that is, the drug-pathway association exists if the target(s) of a drug in $\mathrm{N_{DD}}$ can be mapped on a pathway. The binary associations are weighted based on the count of targets in pathway. The gene-pathway list was integrated from KEGG and UCSC.

**Pathway-pathway association network**

The pathway-pathway association network ($\mathrm{N_{PP}}$) was built based on the hierarchy of KEGG and downloaded from WNS method [8] which integrated the interaction between biological pathways described by KEGG.

#### 4. Parameter selection

In level one model, inter-network jumping probability $\lambda$, global restart probability $\sigma$ together with the contributions of different subnetworks were tuned to optimize the model. The LODOCV method was used as the evaluation strategy and the ROC curves were shown in **Supplementary Figure 1**.

Jumping probability $\lambda_x$ ($\lambda_x \in [0,1]$) refers to the jumping behavior in different subnetworks. The range of $\lambda_x$ is [0,1]. While $\lambda_x = 0$, the jump will be limited in original network (e.g.

$N_{DD}$. On the contrary, if $\lambda_x = 1$, jump will constantly happen between original network and other networks which will lose the ability to discover within networks. For $\lambda \in (0,1)$, larger $\lambda$ means higher probability to jump through inter-network associations (e.g. from $N_{DD}$ to $N_{GG}$ or $N_{PP}$). Jumping probabilities for different subnetworks are not independent ( $\lambda_{DG} = 1 - \lambda_D$, $\lambda_{DP} = 1 - \lambda_D$ and $\lambda_G = \lambda_D$ ), therefore $\lambda_D$ is set to values in $[0, 1]$ for tuning. **Supplementary Figure 1a** shows that changing $\lambda_x$ did not influence much on the performance. AUC values drop while $\lambda = 0$ and $\lambda = 1$ which also reveals the importance of complex drug-gene/pathway network construction. According to tuning result, $\lambda_D = 0.5$ was selected to balance the jumping between networks.

In RWR algorithm, global restart probability $\sigma$ gives the probability that the hypothetical particle can not only jump to the randomly selected neighbor node but also restart in any node in the network. From **Supplementary Figure 1b**, besides $\sigma = 1$ and $\sigma \in [0, 0.3)$, changing $\sigma$ shows trifling effect on results. While $\sigma \in [0, 0.3)$, model suffer from convergence failure. While $\sigma = 1$, the jump of particle in every step is random which results in the AUC of 0.5. The peak of AUC values was about 0.797 when $\sigma = 0.7$ which is consistent with other RWR method based researches [12, 13], thus chosen as final global restart probability.

DrugNet, DGnet, DPnet represent calculating global similarity based on only $N_{DD}$, $N_{DD}$-$N_{DG}$ and $N_{DD}$-$N_{DP}$, respectively. Inter-network jumping probability $\lambda$ and global restart probability $\sigma$ were defined according to tuning described above. Compared with different network construction solutions, combined all sub-networks together shows the best performance (**Supplementary Figure 1c**).

#### 5. Two-threshold strategy for ranking drug pairs

For a drug pair A and B, $r_{A-B}$ is the rank of B when A is used as drug seed and $r_{B-A}$ is the rank of A when B is taken as seed. To avoid the asymmetrical ranks, a two-threshold strategy $(c1, c2)$ was used to integrate $r_{A-B}$ and $r_{B-A}$ and classified drug pairs into combinable, uncombinable and intermediate classes. If both drugs rank before $c1$ or one of the drugs can rank before $c1$ while the other rank falls in intermediate region, the pair will be signed in combinable region. Similarly, if both drugs rank fall after $c2$ or one of the drug ranks in intermediate region while the other fall in the bottom, the outcome will be considered as uncombinable. If both ranks are between $c1$ and $c2$ or one is ranked before $c1$ but the other fall in bottom, the model will consider it as an intermediate pair. We test the possible combination of different thresholds to classify prediction results, and calculate true positive rate (TPR) and true negative rate (TNR). To keep the balance between TPR and TNR, the cutoff "Combinable 20% and uncombinable 50%" were used in level one DComboNet (**Supplementary Figure 2**). For cancer sample specific model, we want to obtain more accurate combinations instead of all potential combinations. Therefore, a stricter cutoff (Top 10% and top 50%) was applied for cancer sample specific model.

#### 6. Experiment validation of drug combination

**Cell culture**

HepG2, SNU475 and SNU423 were purchased from American Type Culture Collection (ATCC, www.atcc.org). Huh7 was purchased from Japanese Collection of Research Bioresources (JCRB, http://cellbank.nibiohn.go.jp/english/). MHCC-97H was gift from

Terence Kin Wah Lee (The University of Hong Kong). CLC5 was established from Chinese liver cancer patient in Lijian Hui' lab. For cell culture, CLC5 was cultured in RPIM 1640 supplemented with 10% FBS, 1 × ITS (Insulin, Transferrin, Selenium Solution) and 40 ng/mL EGF (epithelial growth factor), 100U/ml penicillin and 100μg/ml streptomycin, SNU475 and SNU423 were cultured in RPIM 1640 supplemented with 10% FBS, 100U/ml penicillin and 100μg/ml streptomycin, HepG2, Huh7 and MHCC-97H were cultured in DMEM supplemented with 10% FBS, 100U/ml penicillin and 100μg/ml streptomycin. Both cell lines were maintained in humidified incubator at 37℃ with 5% CO2 and digested by 0.05% trypsin-EDTA for passage at the ratio of 1:3 every 3 days.

**Drug combination screening**

Sorafenib (Cat#S1040), sunitinib (Cat#S1042) and afatinib (Cat#S1011) were purchased from Selleck. The optimal number of seeding cells for each cell line was determined to avoid over-confluency at the end of the drug treatment (around 90% confluency). Cells were seeded in 384-well plates at the pre-determined cell density at a volume of 50 mL by Multidrop Combi Reagent Dispenser (Thermo Fisher Scientific). After overnight incubation, cells were treated with the combination of Sorafenib of 5 or 7 doses and other drugs of 6 doses to yield 5×6 or 7×6 matrix using automated liquid handling platform Bravo (Agilent), and the plates were transferred to the incubator for 72 hr. At the end point of drug treatment, each well was added 25 mL CellTiter-Glo reagent (Promega), and after 10 min incubation in room temperature, the luminescent signals were measured by EnVision Multilabel Reader (PerkinElmer) to determine the cell viabilities. There are four replicate wells for one drug combination dose. The luminescent signal value of each well was normalized to the average value of control wells (DMSO) to calculate the relative cell viability.

### Supplementary figures

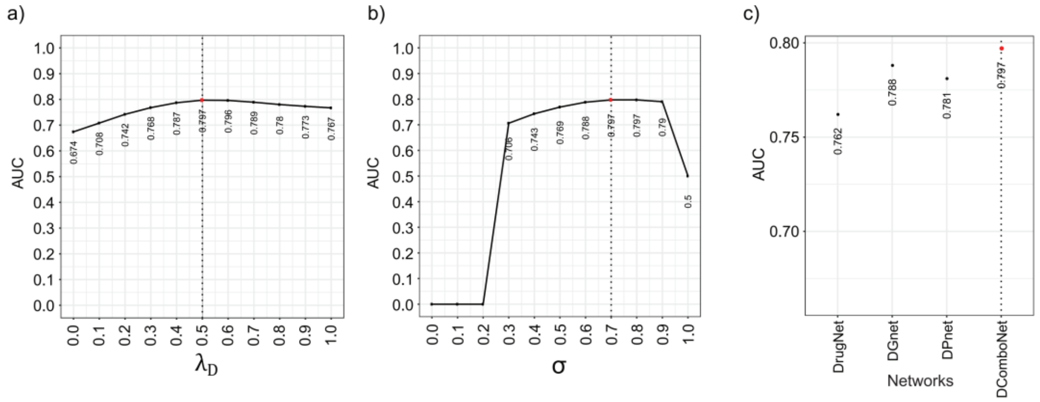

**Figure S1** Parameter selection. a) The influence of jump probability $\lambda_x$, where x-axis denotes the value of corresponding parameter and y-axis represent the AUC values obtained from LOCOCV method, red dots show the selected parameter. b) The influence of global restart probability σ. c) The contribution of different subnetworks on model performance. DComboNet denotes modeling on drug-gene/pathway network, DrugNet which contains only $N_{DD}$, DGnet which contains $N_{DD}$, $N_{DG}$ and $N_{GG}$ and DPnet which contains $N_{DD}$, $N_{DP}$和$N_{PP}$.

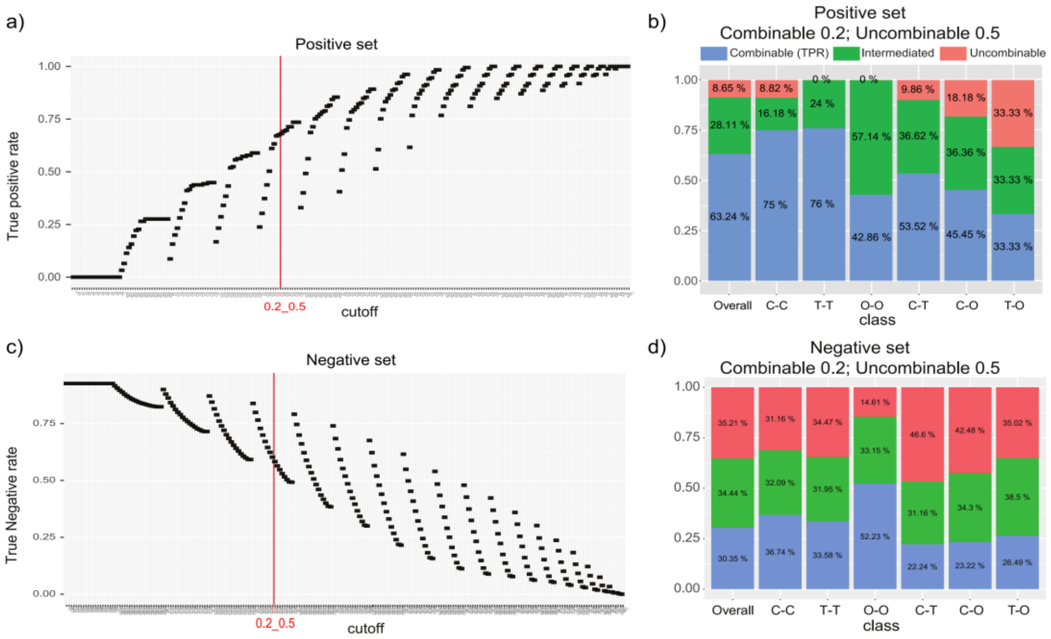

**Figure S2** Two threshold method on DComboNet. a) True positive rate (TPR) on positive set while tuning two threshold method cut-offs. b) The overall TPR and TPRs in different drug combination classes with cut off 'combinable threshold 0.2, uncombinable threshold 0.5', corresponding to **Figure 3b**. c) True negative rate (TNR) on negative set on different tuning two threshold method cut-offs. d) The overall TNR and TNRs in different drug combination classes with cut off 'combinable threshold 0.2, uncombinable threshold 0.5'. a) and c) show with the increasing cut off, TPR increase while TNR drops. From the tuning result, cut off 'combinable threshold 0.2, uncombinable threshold 0.5' (marked in red) shows good balance of prediction power. The color and bar names in b and d) are consistence with **Figure 3b**.

### Supplementary table

**Table S1** Characteristics of known anti-cancer drug combinations

**Table S1A** Features of known anti-cancer drug combinations

**Table S1B** Classification information of drugs in known combinations

**Table S1C** Target genes of drugs in DComboNet

1. Lin, D. *An information-theoretic definition of similarity*. in *Icml*. 1998.
2. Yamanishi, Y., et al., *Drug-target interaction prediction from chemical, genomic and pharmacological data in an integrated framework*. Bioinformatics, 2010. **26**(12): p. i246-54.
3. Yap, C.W., *PaDEL-descriptor: an open source software to calculate molecular descriptors and fingerprints*. J Comput Chem, 2011. **32**(7): p. 1466-74.
4. Bajusz, D., A. Rácz, and K.J.J.o.c. Héberger, *Why is Tanimoto index an appropriate choice for fingerprint-based similarity calculations?* 2015. **7**(1): p. 20.
5. Kuhn, M., et al., *The SIDER database of drugs and side effects*. Nucleic Acids Res, 2016.

**44**(D1): p. D1075-9.
